# The sperm protein SPACA4 is required for efficient fertilization in mice

**DOI:** 10.1101/2021.05.02.442348

**Authors:** Sarah Herberg, Yoshitaka Fujihara, Andreas Blaha, Karin Panser, Kiyonori Kobayashi, Tamara Larasati, Maria Novatchkova, H. Christian Theußl, Olga Olszanska, Masahito Ikawa, Andrea Pauli

## Abstract

Fertilization is the fundamental process that initiates the development of a new individual in all sexually reproducing species. Despite its importance, our understanding of the molecular players that govern mammalian sperm-egg interaction is incomplete, partly because many of the essential factors found in non-mammalian species do not have obvious mammalian homologs. We have recently identified the Ly6/uPAR protein Bouncer as a new, essential fertilization factor in zebrafish (Herberg et al., 2018). Here, we show that Bouncer’s homolog in mammals, SPACA4, is also required for efficient fertilization in mice. In contrast to fish, where Bouncer is expressed specifically in the egg, SPACA4 is expressed exclusively in the sperm. Male knockout mice are severely sub-fertile, and sperm lacking SPACA4 fail to fertilize wild-type eggs *in vitro*. Interestingly, removal of the zona pellucida rescues the fertilization defect of *Spaca4*-deficient sperm *in vitro*, indicating that SPACA4 is not required for the interaction of sperm and the oolemma but rather of sperm and zona pellucida. Our work identifies SPACA4 as an important sperm protein necessary for zona pellucida penetration during mammalian fertilization.

## INTRODUCTION

Fertilization is the fundamental process by which two gametes, the sperm and the egg, fuse to form a single cell which gives rise to a new organism. Despite being essential for all sexually reproducing organisms, the molecular mechanisms that mediate sperm-egg interaction remain poorly understood. One important step towards gaining mechanistic insights into fertilization is the identification of molecules that can mediate sperm-egg interaction. Several proteins that are specifically required for gamete interaction in mammals have been identified with the help of genetic mouse models (reviewed in (Evans, 2012; Krauchunas et al., 2016; Okabe, 2018a)). Required proteins on the sperm are the transmembrane proteins IZUMO1 (Inoue et al., 2005; Satouh et al., 2012), SPACA6 (Lorenzetti et al., 2014; Noda et al., 2020), TMEM95 (Fernandez-Fuertes et al., 2017; Noda et al., 2020; Pausch et al., 2014), FIMP (Fujihara et al., 2020), DCST1/2 (Inoue et al., 2021; Noda et al., 2021) and the secreted protein SOF1 (Noda et al., 2020). The factors required on the egg are the tetraspanin protein CD9 (Kaji et al., 2000; Miyado et al., 2000; Le Naour et al., 2000) as well as the GPI-anchored protein JUNO (Bianchi et al., 2014). The only known interacting protein pair is IZUMO1 and JUNO (Aydin et al., 2016; Bianchi et al., 2014; Han et al., 2016; Kato et al., 2016; Ohto et al., 2016), which are known to mediate adhesion between sperm and egg (Bianchi et al., 2014; Inoue et al., 2005; Kato et al., 2016; Satouh et al., 2012). How the other known mammalian fertilization factors enable sperm-egg binding and/or fusion remains unclear and is subject to active research.

In addition to sperm or egg surface proteins, genetic analyses also revealed the importance of proteins of the mammalian egg coat, called zona pellucida (ZP), in fertilization. ZP proteins were shown to be required for the initial step of sperm binding to the ZP as well as the subsequent block to polyspermy, which is induced by the first sperm entering the egg (Avella et al., 2013, 2014; Gahlay et al., 2010; Rankin et al., 1996, 1998). While IZUMO1, CD9, SPACA6, DCST1/2 and ZP proteins have homologs in non-mammalian vertebrates, phylogenetic analyses by us and others suggest that the other known essential mammalian fertilization factors JUNO, TMEM95, FIMP and SOF1 lack clear non-mammalian homologs (**S1 Fig**) (Grayson, 2015; Pausch et al., 2014). In line with this observation, rapid protein evolution and divergence has been noted as a general hallmark of proteins involved in reproduction (Swanson and Vacquier, 2002).

This evolutionary divergence limits the direct transfer of knowledge gained from studies in invertebrates (Evans and Sherman, 2013; Kosman and Levitan, 2014; Levitan, 2017; Palumbi, 2009) and plants (reviewed in (Dresselhaus et al., 2016)) to fertilization in vertebrates. For example, the mechanistically best-understood fertilization proteins are lysin and Verl from the marine mollusk abalone (reviewed in (Gert and Pauli, 2019)). Lysin is a sperm-expressed and highly abundant secreted protein, whereas Verl is an egg-coat protein that shows structural homology to mammalian ZP2 (Raj et al., 2017). Species-specific binding of lysin to Verl causes non-enzymatic disruption of the abalone egg coat, thereby allowing conspecific sperm to fertilize the egg (Lewis et al., 1982; Lyon and Vacquier, 1999; Raj et al., 2017; Swanson and Vacquier, 1997). However, lysin has no known homolog in vertebrates, leaving it open whether a similar mechanism might contribute to mammalian sperm passage through the ZP.

Apart from the lack of clear homologs, identification of further factors required for sperm-egg interaction in vertebrates has also been hampered by the almost exclusive focus on mammalian fertilization. Mammalian fertilization occurs internally, which poses additional experimental challenges due to the low number of eggs and inaccessibility of gametes. Moreover, possible functional redundancies among fertilization factors have limited the informative value of single-gene knockout studies in mice. Accordingly, many genes that had been implicated as potential fertilization factors based on *in vitro* studies were later shown to be dispensable for fertilization *in vivo* (Ikawa et al., 2010; Okabe, 2018b; Park et al., 2020). Other gamete-specific proteins that might play a role in fertilization have not yet been analyzed for their function *in vivo*, leaving their roles during mammalian fertilization unclear.

One of these gamete-specific proteins is SPACA4 (Sperm Acrosome Associated 4; also called SAMP14 (Sperm Acrosomal Membrane Associated 14)), which was initially identified by mass spectrometry in a screen for membranebound human sperm proteins (Shetty et al., 2003). SPACA4 is particularly interesting for three reasons: First, its fish homolog Bouncer was recently shown to be essential for sperm-egg membrane interaction in zebrafish (Herberg et al., 2018). Secondly, while Bouncer is expressed exclusively in the egg in fish and frogs, its closest mammalian homolog SPACA4 is expressed exclusively in the testis (Herberg et al., 2018; Shetty et al., 2003). Thirdly, the incubation of human sperm with SPACA4-specific antibodies was shown to decrease the binding and fusion of sperm with zona pellucida-free hamster eggs *in vitro* (Shetty et al., 2003). Taken together, these observations point towards an important function of SPACA4 for mammalian fertilization.

Here, we investigate the functional relevance of SPACA4 in mammals by analyzing the phenotypic consequence of genetic loss of SPACA4 in mice.

## RESULTS

### Murine SPACA4 is expressed in testis and localizes to the inner sperm membrane

Bouncer was recently discovered as an essential fertilization factor in zebrafish that is attached to the egg surface via a glycosylphosphatidylinositol (GPI)-anchor and enables sperm binding to the egg membrane (Herberg et al., 2018). Evolutionary analysis revealed that Bouncer has a mammalian homolog, SPACA4 (**Fig 1A**, **Fig S2A**) (Herberg et al., 2018), which raised the immediate question whether SPACA4 might also be important for mammalian reproduction. Bouncer and SPACA4 belong to the large lymphocyte antigen-6 (Ly6)/urokinase-type plasminogen activator receptor (uPAR) protein family, which is characterized by a conserved 60-80 amino acid protein domain containing 8-10 cysteines that adopt a characteristic three-finger fold (Loughner et al., 2016). Most Ly6/uPAR-type genes occur in clusters in the mouse genome (**Fig S2B**), consistent with their origin by gene duplication (Loughner et al., 2016), and are expressed in diverse tissues in mice and humans (**Fig 1B; Fig S2C**). While mammalian *Spaca4* is not the only Ly6/uPAR-type gene expressed specifically in the male germline (testis) (**Fig 1B**; **Fig S2C**), it stands out for having homologs in fish (*bouncer*) and amphibians (*Spaca4*) that are also germline-specifically expressed yet in the opposite sex (ovary) (**Fig 1A**, **Fig S2A**).

**Fig 1.**
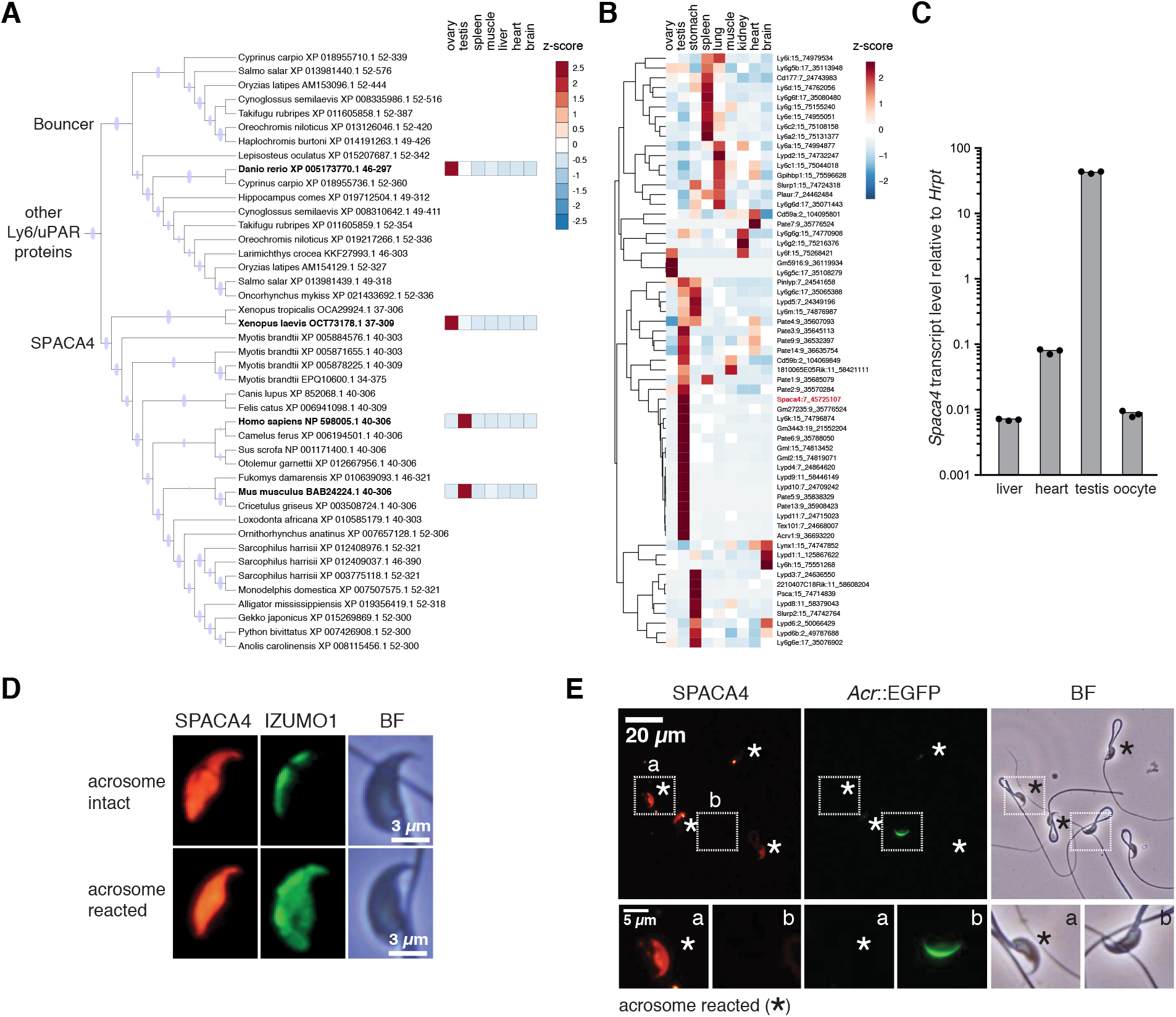
SPACA4 is expressed in murine sperm. (**A**) **Mammalian SPACA4 is the homolog for fish Bouncer and is expressed specifically in testis**. Part of a maximum-likelihood phylogenetic tree based on Ly6/uPAR protein sequence alignments across vertebrates showing SPACA4 and Bouncer (see **Fig S2A** for the full tree; adapted from (Herberg et al., 2018)). Branches supported by ultrafast bootstrap values (>=95%) are marked with a blue dot. Z-scores of averaged gene expression values across adult tissues are shown on the right for *Danio rerio* (Noda et al., 2021), *Xenopus laevis* (Session et al., 2016), *Mus musculus* (Li et al., 2017) and *Homo sapiens* (www.gtexportal.org). While *Bouncer/ Spaca4* mRNAs are expressed in oocytes in fish and frog, mammalian *Spaca4* mRNAs are expressed in testis. (**B**) Mouse Ly6/uPAR genes show diverse expression patterns across adult tissues. The heatmap is color-coded based on z-scores of the normalized gene expression values (average of the square-root) of RNA-Seq data from murine adult tissues (Li et al., 2017). The clustering and dendrogram (left) are based on expression scores. *Spaca4* is highlighted in red. Numbers behind gene names indicate chromosome location in mice. (**C**) **Mouse Spaca4 RNA is expressed in the male germline**. RT-qPCR from murine cDNA from different adult tissues reveals enrichment of *Spaca4* specifically in mouse testis. Primers amplifying *Hypoxanthineguanine phosphoribosyl transferase* (*Hrpt*) were used as a control. (**D**) **Mouse SPACA4 protein localizes to the sperm head**. Immunostaining of SPACA4 (red) and IZUMO1 (green) under permeabilizing conditions detects SPACA4 and IZUMO1 in the sperm head. In contrast to IZUMO1 that relocalizes after the acrosome reaction, SPACA4 does not change its localization. BF, brightfield image. Scale bar, 3 μm. (**E**) **Murine SPACA4 localizes to the inner sperm membrane**. Immunofluorescence staining of SPACA4 (red) in mouse spermatozoa under non-permeabilizing conditions. Spermatozoa were derived from transgenic mice expressing EGFP under the control of the Acrosin promoter (Nakanishi et al., 1999), labeling sperm with intact acrosomes (green). Acrosome-reacted spermatozoa are highlighted by an asterisk. Boxed areas are shown below at higher magnification (a – acrosome-reacted sperm; b – spermatozoa with intact acrosome). SPACA4 was detected in acrosome-reacted spermatozoa (a) but not in spermatozoa with intact acrosomes (b). BF, brightfield image. Scale bar, 20 μm (top) and 5 μm (bottom).

To confirm that *Spaca4* was indeed expressed in male but not female gametes in mice, we analyzed the expression level of *Spaca4* mRNA in different tissues using RT-qPCR. Murine *Spaca4* was detected specifically in testis and enriched 100-to 1000-fold compared to other tissues (**Fig 1C**; **Fig S2D**), which agrees with published RNA-Seq data from mice (**Fig 1A,B**) and with the reported testis-specific expression in humans (Shetty et al., 2003). Analysis of published singlecell RNA-Seq data from murine spermatogenesis revealed a peak of *Spaca4* mRNA expression in round spermatids, which resembles the expression of *Izumo1* mRNA in timing and magnitude (**Fig S3**) (Ernst et al., 2019). Using anti-mouse SPACA4 antibodies, SPACA4 was found to localize to the sperm head (**Fig 1D**) and was readily detected on acrosome-reacted but not on acrosome-intact live sperm (**Fig 1E**). This expression pattern is consistent with the reported acrosomal membrane localization of human SPACA4 (Shetty et al., 2003).

### Male mice lacking SPACA4 are sub-fertile

To investigate the function of mammalian SPACA4 *in vivo*, we generated *Spaca4* knockout mice by CRISPR/Cas9-mediated targeted mutagenesis. We recovered two mutant alleles in the C57BL/6J background. The first one, in the following called *Spaca4^77del^*, contains a 77-nt deletion leading to a frameshift after amino acid 42 (**Fig 2A-B**; **Fig S4A**). The second allele, in the following called *Spaca4^117del^*, contains a 117-nt inframe deletion, which removes half (39 amino acids) of the mature Spaca4 protein and is thus also predicted to result in a full knockout mutation (**Fig 2A-B**; **Fig S4A**). An independent *Spaca4* knockout mouse (*Spaca4*^tm1Osb^) was generated in the BDF1 background by replacing the whole exon of the *Spaca4* gene with a neomycin resistance cassette using homologous recombination (**Fig S5A-C**). *Spaca4* knockout mice appeared indistinguishable from wild-type and heterozygous littermates, revealing that SPACA4 is dispensable for somatic development in mice.

**Fig 2.**
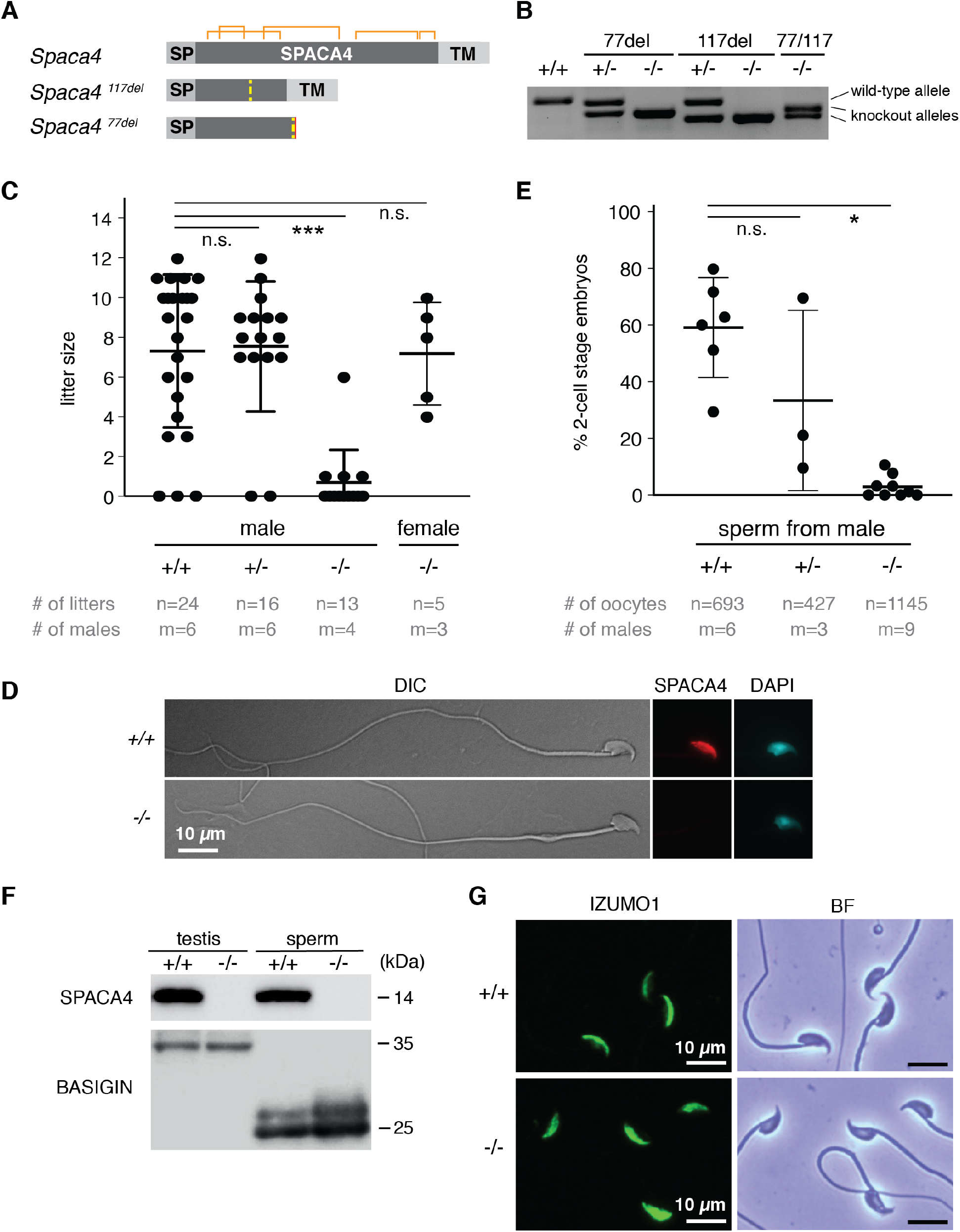
SPACA4 is required for efficient fertilization in male mice. (**A** and **B**) **Overview of the C57BL/6J-*Spaca4* knockout alleles generated by CRISPR/Cas9-mediated targeted mutagenesis**. One allele contains a 117-nt in-frame deletion after amino acid 42. The other allele contains a 77-nt out-of-frame deletion after amino acid 47. (**A**) Schematic of the wild-type and knockout alleles. Yellow dashed lines indicate the site of the deletions. Predicted disulfide bridges are indicated in orange. SP, signal peptide; TM, transmembrane region. (**B**) Genotyping of *Spaca4* knockout mice. *Spaca4* PCR products are separated based on their sizes on an Agarose-gel. (**C**) ***Spaca4* knockout male mice are sub-fertile**. Litter sizes of C57BL/6J-*Spaca4* wild-type (+/+), heterozygous (+/−) and transheterozygous (−/−) males caged with B6129F1 wild-type females, or B6129F1 wild-type males caged with transheterozygous (−/−) females. Successful mating was confirmed by plug checks. Data are means ± SD. ***p < 0.0001 (Kruskal-Wallis test with Dunn multiple-comparisons test); n.s., not significant. n = number of litters; m = number of male mice tested. (**D**) **Sperm morphology is normal in the absence of SPACA4 protein**. Immunostaining of sperm detects SPACA4 protein (red) under permeabilizing conditions in the head of sperm from wild-type (+/+) mice but not in sperm from *Spaca4* knockout (−/−) mice. Sperm morphology is normal. DAPI (cyan) staining labels the sperm nucleus. DIC, differential interference contrast image. Scale bar, 10 μm. (**E**) **Sperm from *Spaca4* knockout mice has a severely reduced ability to fertilize wild-type oocytes**. *In vitro* fertilization of oocytes from C57BL/6J wild-type females using sperm from C57BL/6J-*Spaca4* wild-type (+/+), heterozygous (+/−) or transheterozygous (−/−) males. Plotted is the percentage of 2-cell stage embryos as a measure of successful fertilization. Data are means ± SD. *p = 0.014 (Kruskal-Wallis test with Dunn multiplecomparisons test); n.s., not significant. n = total number of oocytes; m = number of males tested. (**F**) **SPACA4 protein is absent in sperm and testis from *Spaca4* knockout mice**. Western blot analysis showing the absence of SPACA4 protein in testes and spermatozoa of *Spaca4* knockout mice. Testes and spermatozoa of wild-type mice as well as BASIGIN protein, which undergoes proteolytic cleavage during sperm maturation (Noda et al., 2020), are used as control. (**G**) **IZUMO1 localization is normal in *Spaca4* knockout mice**. Immunostaining of sperm shows normal localization of IZUMO1 (green) in wild-type and SPACA4-deficient sperm. BF, brightfield image. Scale bar, 10 μm.

In line with SPACA4’s sperm-specific expression in mammals, we found that SPACA4 is necessary for male fertility. The litter size of transheterozygous (*Spaca4^117del/77del^*) as well as homozygous mutant male mice was significantly lower than the litter sizes of wild-type or heterozygous male mice (p < 0.0001) (**Fig 2C**; **Fig S4B and S5D**). In contrast, female fertility was not affected by the *Spaca4* mutation (**Fig 2C**). The observed defect in male fertility was not due to an inability of *Spaca4* knockout males to mate as verified by the presence of vaginal plugs.

The severely reduced fertility of males lacking SPACA4 could have multiple reasons, including reduced sperm count, immotility, or a defect in gamete interaction. Sperm morphology, numbers and overall sperm motility were similar in wildtype and knockout mice (**Fig 2D**, **Fig S4C-E**), suggesting that SPACA4 is required during sperm-egg interaction. To test this hypothesis, we performed *in vitro* fertilization (IVF) experiments, which revealed that *Spaca4* mutant sperm was severely compromised in its ability to fertilize wild-type oocytes *in vitro*: Spermatozoa from transheterozygous (*Spaca4*^*117*del/*77*del^) male mice resulted in a severely reduced average fertilization rate of 2.9% (2-cell stage embryos), whereas the average fertilization rates using spermatozoa from wild-type or heterozygous (*Spaca4*^*117*del/+^ or *Spaca4^77del/+^*) mice were at 59.4% or 33.4%, respectively (**Fig 2E**). Similar defects in IVF were observed in *Spaca4*^tm1Osb−/−^ sperm (**Fig S5E**). The inability of SPACA4-deficient sperm to fertilize wild-type oocytes could be due to loss of expression and/or improper localization of IZUMO1. However, Western blotting and immunofluorescence staining revealed that sperm of *Spaca4* knockout mice showed normal expression of IZUMO1 (**Fig 2F-G**). Overall, we conclude that SPACA4 is an important albeit not absolutely essential protein for mammalian gamete interaction.

### SPACA4 is required for efficient penetration of the zona pellucida

The marked *in vitro* fertilization defect of sperm derived from *Spaca4*^−/−^ males showed that SPACA4 is necessary for sperm to efficiently interact with the egg. In mammals, this interaction occurs in two steps: sperm first needs to bind and penetrate the zona pellucida (ZP) before binding to the egg membrane (oolemma), which enables sperm-egg fusion. To determine at which stage of fertilization SPACA4-deficient sperm is impaired, we quantified the number of spermatozoa bound to the ZP 30 minutes after insemination. We found that in both *Spaca4*^−/−^ mutant strains, fewer mutant spermatozoa bound the ZP of wild-type oocytes compared to control wild-type sperm (23.5 ± 9.5 in the case of *Spaca4^117/77^* vs. 30.7 ± 7.0 for wild type (BJ6) (p < 0.001); 34.3 ± 11.1 in the case of *Spaca4*^tm1Osb^ vs. 59.7 ± 10.7 for wild type (BDF1) (p < 0.001)) (**Fig 3A**). The reduced binding of SPACA4-deficient sperm to the ZP, combined with a reduced motility of mutant sperm during incubation in IVF medium (**Fig S6**), prompted us to ask whether SPACA4 might be involved in the penetration of the protective layers that surround the oocyte, namely the cumulus layer and the ZP. To test whether removal of the cumulus cells and/or the ZP can rescue the fertility defect of SPACA4-deficient sperm in IVF, wild-type oocytes were treated with hyaluronidase and acidified Tyrode’s solution to remove the cumulus layer and ZP, respectively (Richard Behringer, Marina Gertsenstein, Kristina Nagy, 2014) (**Fig 3B**). Fertilization by wild-type sperm was not significantly affected by the different treatments (**Fig 3C**). Similarly, removal of the cumulus cells did not rescue the low fertilization rate of SPACA4-deficient sperm (18.8% (COCs) vs. 4.7% (cumulus cells removed) (p = 0.884); **Fig 3C**). However, removal of the ZP enabled SPACA4-deficient sperm to fertilize zona-free oocytes at a similarly high rate (95.8%) as wild-type sperm (97.1%) (p-value for mutant sperm fertilizing oocytes with ZP vs. ZP-free = 0.033) (**Fig 3C**). We therefore conclude that murine SPACA4 is required for the sperm’s ability to efficiently traverse the ZP (**Fig 3D**).

**Fig 3.**
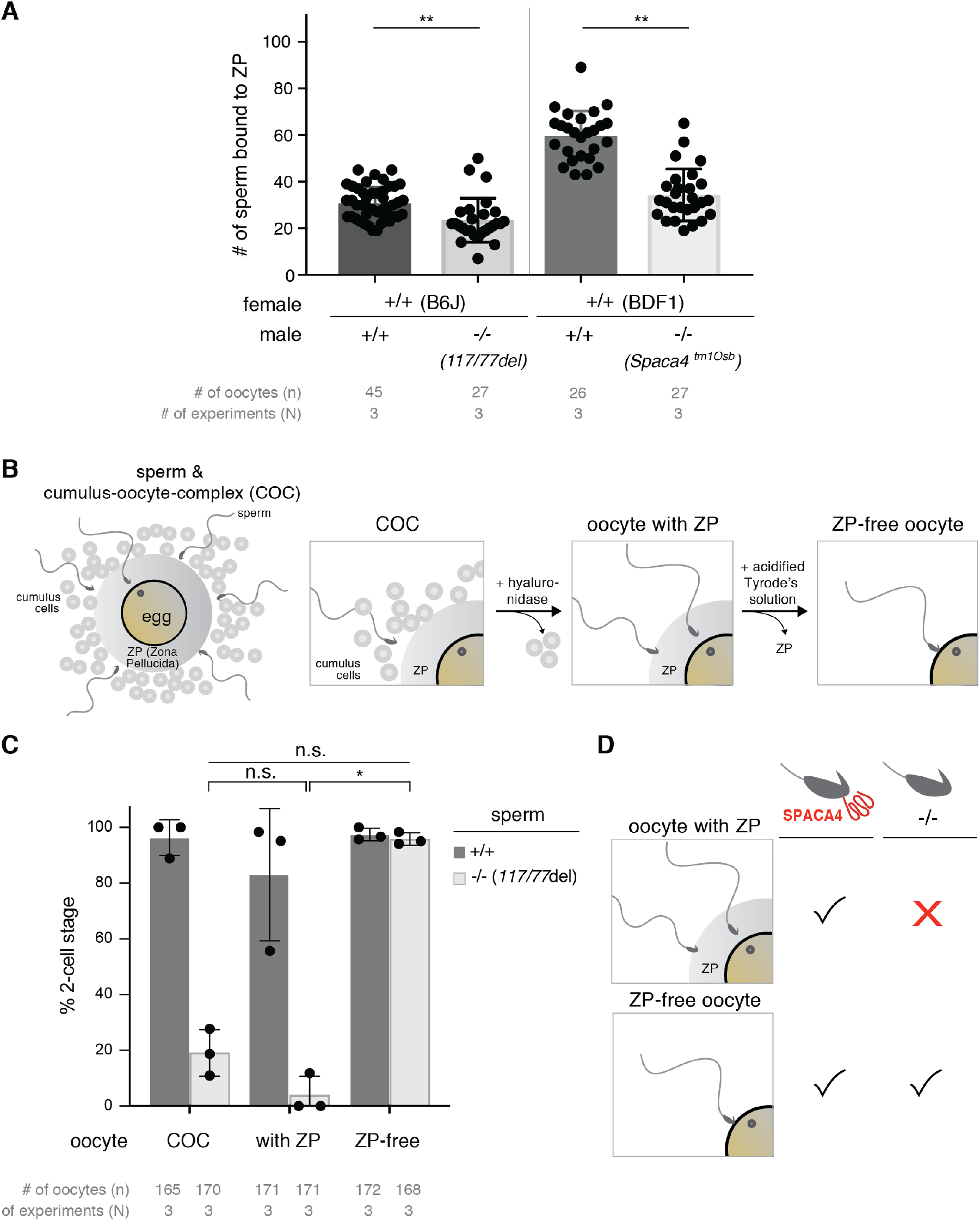
SPACA4 enables sperm to bind to and penetrate the zona pellucida. (**A**) **SPACA4 is required for efficient binding of sperm to the ZP.** Sperm of the indicated genotypes was incubated for 30 minutes with THY-treated oocytes (with intact ZP) of matching genetic backgrounds. Plotted is the number of sperm bound to ZP-containing oocytes. Data are means ± SD. P-values are of Student’s t test. n = total number of oocytes; N = number of replicates. (**B**) **Schematic of sperm bound to the cumulus-oocyte-complex (COC)**, and experimental treatments used to remove the cumulus cells (by treatment of COCs with hyaluronidase) and the zona pellucida (ZP) (by treatment of cumulus-free oocytes with acidified Tyrode’s solution) from the COCs. (**C**) **SPACA4 is required for ZP penetration but not for oolemma binding and fusion.** IVF performed with COCs from super-ovulated C57BL/6J females, cumulus cell-free oocytes (oocyte with ZP), and ZP-free oocytes with sperm from either wild-type C57BL/6J males or age-matched *Spaca4^117nt-del/77nt-del^* males. Plotted is the percentage of 2-cell stage embryos as a measure of successful fertilization. Data are means ± SD. P-values are of a Kruskal-Wallis test with Dunn multiple-comparisons; n.s., not significant. n = total number of oocytes; N = number of replicates. (**D**) **Model of SPACA4’s function in murine fertilization.** SPACA4 is required for efficient ZP penetration in mice.

## DISCUSSION

Here we reveal that SPACA4, the mammalian homolog of fish Bouncer (Herberg et al., 2018), is required for normal male fertility in mice (**Fig 2**). We find that SPACA4 is expressed in the sperm head and gets exposed by the acrosome reaction (**Fig. 1D-E**), which enables sperm to efficiently bind to and penetrate the zona pellucida (**Fig 3**). Thus, our work provides genetic evidence for an important function of SPACA4 for reproduction in mice. Together with previous SPACA4-antibody blocking experiments with human sperm and the conserved expression pattern of human SPACA4 in sperm (Shetty et al., 2003), our results in mice could have direct relevance for male fertility in humans. Future work will be needed to explore a possible link between *Spaca4* mutations in humans and sub-fertility in men presenting normal sperm count, morphology and motility.

SPACA4 and Bouncer are not the only Ly6/ uPAR proteins that have been linked to vertebrate reproduction. Several other members of this large gene family show a testis-restricted expression pattern in mammals (**Fig 1B**, **Fig S2C**), some of which (*Tex101, Lypd4, Pate4* and the *Pate* gene cluster) have been confirmed in genetic knockout studies to be required for male fertility in mice (Fujihara et al., 2014, 2019; Noda et al., 2019). In light of these known male-specific requirements for Ly6/uPAR proteins in mammals, SPACA4 and Bouncer present an interesting example of homologous proteins that diverged in terms of gene expression pattern and function. Our phylogenetic sequence analysis shows that Bouncer and SPACA4 are the closest homologs among all other Ly6/uPAR family members (**Fig S2A)**, yet they have opposing germline-specific expression patterns that broadly correlate with external (fish, amphibians; expressed in the egg) versus internal (mammals, reptiles; expressed in the sperm) fertilization (**Fig 1A, Fig S2A).** We currently do not know how this different gene expression pattern arose, and whether it evolved as a consequence of the different fertilization modes. One possibility is that an ancestral Ly6/uPAR protein might have been expressed in both male and female gonads, and that sex-specific loss of expression occurred either by chance in different lineages or in response to a functional benefit of the restricted expression of SPACA4/Bouncer to the male or female germline. Acquisition of a restricted expression domain from an initially broader expression pattern has been proposed for other members of the Ly6/uPAR protein family, namely for snake toxins that evolved a venom gland-restricted expression pattern (Hargreaves et al., 2014).

The difference in expression pattern also extends to a difference in functionality at least in the case of the two example model organisms zebrafish (Herberg et al., 2018) and mice (this work): While Bouncer in fish is required in the egg for sperm binding to the oolemma (Herberg et al., 2018), the results presented here reveal that SPACA4 is dispensable for sperm binding to the oolemma in mice and instead required for the preceding process of zona pellucida penetration and/or binding. In this regard it is interesting to note that mammalian and fish gametes differ in key aspects: Mammalian sperm has an acrosome, a specialized vesicle in the sperm head that must undergo exocytosis to expose important membrane-localized fertility factors to become exposed on the surface of the sperm head (Hirohashi and Yanagimachi, 2018; Satouh et al., 2012). This so-called acrosome reaction is important for successful zona pellucida penetration and fertilization. Moreover, mammalian sperm needs to first bind to and penetrate the outer coat before gaining access to the egg membrane. Fish sperm, on the other hand, lacks an acrosome and has direct access to the oolemma through the micropyle, a pre-formed funnel in the outer protective layer of the fish egg. One can therefore speculate that acquisition of a sperm-specific expression of SPACA4 in mammals was beneficial to allow sperm passing the additional outer barrier. In zebrafish, current data suggests that Bouncer acts in a unilateral manner by interacting with a still unknown factor on the opposing gamete since (1) expression in sperm does not rescue infertility of *bouncer* mutant females, and (2) successful sperm entry requires compatibility between a species’ Bouncer and the sperm (Herberg et al., 2018). Whether a similar mode of action, e.g. a SPACA4 interacting protein expressed in the oocyte and a possible involvement in species-specificity of sperm-egg interaction, also applies to mammals is currently unclear. Given the divergence in function of Ly6/uPAR proteins (Bychkov et al., 2017; Loughner et al., 2016), it is possible that SPACA4 acquired a different mode of action from Bouncer, e.g. by acting *in cis* through interacting with other sperm-expressed membrane proteins, or by interacting with oviductal proteins as suggested for its interaction with plasminogen (Ferrer et al., 2016). Moreover, one can speculate that for species with internal fertilization, a selection step determining gamete compatibility at the stage of sperm-egg interaction might be less important since mating partner selection alone can ensure that only the selected partner’s sperm will be available for fertilization. This is not the case for species performing external fertilization who cannot guarantee by pre-mating choice that only conspecific sperm reaches the egg (Levitan, 2017; Levitan and Ferrell, 2006), and whose oocyte-specific expression of Bouncer could contribute to post-copulation female mate choice (also called cryptic female mate choice) (Firman et al., 2017). Thus, sperm-expressed proteins like SPACA4 could promote the efficiency of fertilization while oocyte-expressed proteins like Bouncer could support the selection of conspecific sperm. Future experiments will be required to elucidate how SPACA4 promotes fertilization in mammals and to which extent the mechanism differs between SPACA4 and Bouncer.

Overall, our study on SPACA4/Bouncer highlights an interesting example of a vertebrate-specific sperm-egg interaction protein that evolved a different function and gene expression pattern in fish versus mammals.

## ACKNOWLEDGMENTS

We thank the animal facility personnel at IMP/ IMBA, in particular Jasmin Ecker and Lukas Laister, and Naoko Nagasawa at Osaka University/ National Cerebral and Cardiovascular Center for excellent care of animals and for help with mouse husbandry; J.R. Wojciechowski, L. Haas from the Obenauf lab, A. Fedl from the Busslinger lab, D. Keays and I. Baptista from the Keays lab, and E. Chatzidaki and J. Gassler from the Tachibana lab for technical assistance and advice with mouse work; K. Aumayer and her team of the biooptics facility at the ViennaBiocenter for support with microscopy; the entire Pauli lab, particularly Krista Gert and Victoria Deneke, for valuable discussions and comments on the project and manuscript.

## AUTHOR CONTRIBUTIONS

Y.F., A.P and M.I. designed the study. S.H., Y.F., A.B., K.P., K.K. and T.L. performed experiments with the help of K.K., T.L., H.C.T. and O.O. S.H., Y.F., A.B., K.P., K.K., T.L., H.C.T., M.I. and A.P. analyzed the data. M.N. performed evolutionary and gene expression analyses. A.P. wrote the manuscript with help of S.H., Y.F., A.B. and M.I. and input from all authors.

## MATERIALS AND METHODS

### Mouse lines and husbandry

All mouse experiments were conducted according to Austrian and European guidelines for animal research and approved by local Austrian authorities or by the Animal Care and Use Committee of the Research Institute for Microbial Diseases, Osaka University, Japan (#Biken-AP-H30-01). Mice were maintained under a 10/14-hr light/dark cycle (IMP, Vienna, Austria) or under a 12-hr light/dark cycle (Osaka University, Japan). Wild-type mice were purchased from CLEA Japan (Tokyo, Japan) and Japan SLC (Shizuoka, Japan) (Osaka University, Japan). The mouse strain Tg(Acr-EGFP)1Osb has been reported before (Nakanishi et al., 1999). The *Spaca4* knockout mice, C57BL/6N-*Spaca4*^tm1Osb^, were deposited to the RIKEN BioResource Center (http://mus.brc.riken.jp/en/) and are available to the scientific community.

### Generation of *Spaca4* knockout mice

The mouse *Spaca4* gene consists of a single exon and maps to chromosome 7. Two strategies were used to generate *Spaca4* knockout mice: 1) CRISPR-Cas9-based gene targeting (IMP, Vienna, Austria); and 2) replacement of the Spaca4-encoding exon with a neomycin selection cassette via gene targeting of ES-cells (Osaka University, Japan).

#### CRISPR-Cas9-based gene targeting

(resultant *Spaca4* alleles: C57BL/6J-*Spaca4*^*117*del^ and C57BL/ 6J-Spaca4^*77del*^): The *Spaca4* knockout mice were generated using CRISPR-Cas9-based gene targeting. Two guide RNAs (gRNAs) targeting the coding region of *Spaca4* were generated according to published protocols (Gagnon et al., 2014) by oligo annealing followed by T7 polymerase-driven *in vitro* transcription (gene-specific targeting oligo: SPACA4_gRNA1 and 2; common tracer oligo). For gRNA injections, zygotes were isolated from super-ovulated donor female (C57BL/6J) mice on the day of the coagulation plug (= E0.5). To remove cumulus cells, zygotes were incubated in hyaluronidase solution (~0.3 mg/ml). The injection mix (50 ng/μl *Cas9* mRNA (Sigma-Aldrich), 50 ng/μl SPACA4_gRNA1 and 2) was injected into the cytoplasm of the zygotes. Injected zygotes were incubated for at least 15 minutes at 37°C and 5% CO2. Surviving zygotes were transferred into the oviducts of pseudo-pregnant recipient females. The resulting pubs were genotyped using primers SPACA4_gt_F and SPACA4_gt_R.

#### Gene targeting in ES cells

(resultant *Spaca4* allele: C57BL/6N-*Spaca4*^tm1Osb^) The targeting vector was constructed using pNT1.1 (https://www.ncbi.nlm.nih.gov/nuccore/JN935771). A 1.9 kb NotI-XhoI short arm fragment and a 5.1 kb PacI-MfeI long arm fragment were obtained by PCR amplification using BAC DNA (RP24-343L2) as a template. The primers used were: Spaca4_targeting-s_F, Spaca4_targeting-s_R, for the short arm; Spaca4_targeting-l_F, Spaca4_targeting-l_R for the long arm. The targeting construct was linearized with ClaI and electroporated into EGR-G101 [C57BL/6N-*Tg*(*CAG/Acr-EGFP*)] embryonic stem (ES) cells (Fujihara et al., 2013a). Potentially targeted ES cell clones were separated by positive/negative selection with G418 and ganciclovir. Correct targeting of the *Spaca4* allele in ES cell clones and germline transmission were determined by PCR. Screening primer sused were: Spaca 4_screening + gt#781 and Spaca4_screening+gt#5081 for the short arm; Spaca 4_screening + gt#5 173 and Spaca4_screening+gt#678 for the long arm. The mutant ES clones were injected into 8-cell stage ICR embryos, and chimeric blastocysts were transferred into the uterine horns of pseudopregnant ICR females the next day. To confirm germline transmission, chimeric males were mated with B6D2F1 females. Offspring from heterozygous intercrosses were genotyped by PCR. The genotyping primers used were: Spaca4_gt#5269, Spaca4_gt#5298, and Spaca4_screening+gt#781. Two bands, a 0.3-kb band as the wild-type allele and a 0.5-kb band as the knockout allele, were amplified by PCR.

### Genotyping: Extraction of gDNA from mouse ear clips and genotyping PCR

Mice were genotyped at weaning age (around 3 weeks) using ear clips. gDNA was extracted from the ear clips using one of the following three alternative protocols. According to one protocol, ear clips were lysed by incubation in 25 μl of QuickExtract Solution (QE09050, Lucigen) at 65 °C and shaking at 600 rpm for 30 min. According to a second protocol, ear clips were lysed in 100 μl lysis buffer (0.1 M NaCl, 0.5% SDS, 10 mM TRIS pH 8.0, 0.25 mM EDTA, 2μg/μl proteinase K) at 55 °C for 3 hours. To precipitate the cellular debris 60 μl NaCl were added and the sample was centrifuged at (21000 g) for 10 min at 4 °C. The supernatant was centrifuged a second time (21000 g, 10 min at 4 °C). To precipitate the DNA, 160 μl of cold 100% ethanol was added, and the sample was centrifuged again (21000 g, 10 min at 4 °C). The pellet was washed a second time using 250 μl of 75% ethanol (centrifugation at 21000 g, 5 min at 4 °C). The cleaned DNA pellet was dried at 37 °C for 5-15 min and solubilized in 50 μl ddH2O. According to a third protocol, a commercial lysis buffer was used to extract the gDNA from ear clippings (DirectPCR Lysis Reagent (Mouse Tail), ViagenBiotech). 200 μl of the commercial lysis buffer and 1 μl of 100 mg/ml proteinase K were added to each ear clip and incubated at 55 °C under shaking (600 rpm) overnight. For heat inactivation the samples were heated to 85 °C for 45 min under vigorous shaking (800 rpm).

To amplify the *Spaca4* coding region, standard Taq Polyermase or Q5 Hot Start polymerase (New England Biolabs) was used with primers SPACA4_gt_F and SPACA4_gt_R according to the manufacturer’s protocol. The size of the PCR product was analyzed on a 2% agarose gel.

### Identification of the nature of the *SPACA4^77del^* and *SPACA4^117del^* mutations

To analyze the nature of the *Spaca4* mutations, the PCR products from the genotyping of the first outcross (generation F1) were cloned into the cloning vectors provided by the StrataClone PCR cloning kit (Stratagene). The cloning was performed according to the manufacturer’s protocol. Of the resulting bacterial colonies 96 colonies were picked and sequenced using primer SPACA4_gt_F. Two mice, one with a 77-nt and the other one with a 117-nt deletion in the coding sequence of *Spaca4* were selected (see below) and used for further in- and out-crossing. Both deletions were also confirmed by sequencing the PCR products from the genotyping of the next generation (F2).

#### Wild-type *Spaca4* ORF

Atggtccttggctggccactgcttctggtgttggttctttgcccaggtgtgacaggcatcaaggactgcgtcttctgtgagctgactgactctgctcggtgccctggcacacacatgcgctgt<u>ggggatgacgaagattgctt**cacaggccacggagtagcccagggtgtggggcccatcatcaacaaaggctgcgtgcactccaccagctgtggccgcg**aggaacccatcagctacatg</u>ggcctcacatacagtctcaccaccacctgctgttctggccacctttgcaataagggcactggcctttccacaggggctaccagcctgtcactgggtctgcagctgctcctgggcctgttgctgctgcttcaatactggctgtga

##### Bold sequence

is deleted in the mutant *Spaca4^77del^ <u>Underlined sequence</u>* is deleted in the mutant *Spaca4^117del^*

#### Wild-type SPACA4 protein

<u>MVLGWPLLLVLVLCPGVTG</u>IKDCVFCELTDSARC *PGTHMRCG<u>DDEDCFTGHGVAQGVGPIINKGCVHST</u>SCGREEPISYMG*LTYSLTTTCCSGHLCNKGTGLSTG ATSLSLGLQLLLGLLLLLQYWL*

Underlined sequence: signal peptide

##### Bold sequence

is deleted in the mutant *Spaca4^77del^*

<u>Underlined italics sequence</u> is deleted in the mutant Spaca4^117del^

#### *Spaca477del* ORF

*Atggtccttggctggccactgcttctggtgttggttctttgcccaggtgtgacaggcatcaaggactgcgtcttctgtgagctgactgactctgctcggtgccctggcacacacatgcgctgtggggatgacgaagattgcttaggaacccatcagctacatgggcctcacatacagtctcaccaccacctgctgttctggccacctttgcaataa*

#### SPACA4^77del^ protein

<u>MVLGWPLLLVLVLCPGVTG</u>IKDCVFCELTDSARC PGTHMRCGDDEDC**LGTHQLHGPHIQSHHHLLF WPPLQ\***

(**bold sequence** does not exist in the wild type as it is a consequence of the out-of-frame deletion.)

##### *Spaca4^117del^* ORF

*Atggtccttggctggccactgcttctggtgttggttctttgcccaggtgtgacaggcatcaaggactgcgtcttctgtgagctgactgactctgctcggtgccctggcacacacatgcgctgtggcctcacatacagtctcaccaccacctgctgttctggccacctttgcaataagggcactggcctttccacaggggctaccagcctgtcactgggtctgcagctgctcctgggcctgttgctgctgcttcaatactggctgtga*

#### SPACA4^117del^ protein

<u>MVLGWPLLLVLVLCPGVTG</u>IKDCVFCELTDSARC PGTHMRCGLTYSLTTTCCSGHLCNKGTGLSTGAT SLSLGLQLLLGLLLLLQYWL*

### RT-PCR and RT-qPCR of *Spaca4*

Mouse cDNA was prepared from multiple adult tissues of ICR mice and from testes of *Spaca4* knockout mice. Total RNA was reverse-transcribed into cDNA using a SuperScript III First-Strand Synthesis System for RT-PCR (Invitrogen). The amplification conditions for PCR were 2 min at 50°C, 30 sec at 95°C, followed by 39 cycles of 95°C for 15 sec and 60°C for 1 min (+ plate read) (RT-qPCR **Fig 1C**) or 1 min at 94°C, followed by 30 cycles of 94°C for 30 sec, 65°C for 30 sec, and 72°C for 30 sec, with a final 7 min extension at 72°C (RT-PCR **Fig S2D**), using primers targeting *Spaca4* (Spaca4_qPCR_F2 and Spaca4_qPCR_R2) and *Hprt* (HPRT_qPCR_F and HPRT_qPCR_R) (**Fig 1C**) or *Spaca4* (Spaca4_qPCR_F1 and Spaca4_qPCR_R1) and *Gapdh* (Gapdh_RT_F and Gapdh_RT_R) (**S2D Fig**).

### *In vivo* fertility assays

Fertility assays in mice were performed according to two alternative methods: In the case of CRISPR-generated C57BL/6J-*Spaca4* alleles, C57BL/6J-*Spaca4* wild-type, heterozygous, transheterozygous (*Spaca4*^*117*del/77*del*^) or homozygous (*Spaca4*^*117*del/117del^ or *Spaca4*^*77*del/77*del*^) mutant male or female mice were caged with 2-4-month-old B6129F1 wild-type mice in the evening. Females were checked for plugs every morning and separated from the males as soon as a plug could be observed. The number of pups for each female was counted within a week of birth. In case of the mutant male mice, this procedure was repeated at least once before the mutant mice were kept caged with a B6129F1 female for 3-10 weeks after the initial plug.

In the case of C57BL/6N-*Spaca4*^tm1Osb^ mutants, sexually mature wild-type, heterozygous or homozygous mutant male mice were caged with 2-month-old B6D2F1 for several months, and the number of pups in each cage was counted within a week of birth. Average litter sizes are presented as the number of total pups born divided by the number of litters for each genotype.

### *In vitro* fertilization assays

Before *in vitro* fertilization, female mice were superovulated by injection of CARD HyperOva (KYD-010-EX, Cosmo Bio Co) approx. 63 hours before and hCG (Chorulon) 14-16 hours before harvesting the oocytes. *In vitro* fertilization was performed using CARD MEDIUM (KYD-003-EX, Cosmo Bio Co) and CARD FERTIUP Preincubation Medium (KYD-002-EX, Cosmo Bio Co) according to the manufacturer’s protocol. Sperm was prepared from the cauda epididymides and capacitated in CARD FERTIUP medium for 1 hour. Oocytes from superovulated female mice (C57BL/6J) were introduced into a drop of CARD MEDIUM. To prepare cumulus- or zona-free oocytes (**Fig 3C**), cumulus-oocyte-complexes (COCs) were collected in M2 medium (MR-015P-5D, Merck) and treated with 300 μg/ mL hyaluronidase (H3884, Sigma-Aldrich) until the cumulus cells were removed, and washed in M2. For ZP-removal, the cumulus cell-free oocytes were moved to a droplet of acidified Tyrode’s solution (T1788, Sigma-Aldrich) for a few seconds, then washed with M2 and finally transferred into CARD MEDIUM. Afterwards, the preincubated sperm was added to the differently treated oocytes for fertilization. Sperm and eggs were incubated at 37°C and 5% CO2 and washed 3 hours after incubation. Fertilization rates were recorded by counting the number of two-cell stage embryos on the next day.

### Sperm number and motility analyses

The number (**Fig S4D**) and overall motility (motile, progressive motile; **Fig S4E**) of sperm were measured using the computer-assisted sperm analysis system CEROS II animal (Hamilton Thorne) according to the manufacturer’s protocol. In brief, 1μl of sperm was diluted 1:200 in DPBS (MR-006C, Merck) and then overall motility assessed on a CEROS II.

To quantify sperm motility under IVF conditions **(Fig S6**), cauda epididymal spermatozoa were squeezed out, and then dispersed in TYH (for sperm motility and IVF). After incubation of 10 and 120 minutes in TYH, sperm motility patterns were examined using the CEROS II sperm analysis system (Goodson et al., 2011; Noda et al., 2019).

### Assessment of sperm binding to the zona pellucida

Sperm ZP-binding assay was performed as described previously (Yamaguchi et al., 2006). Briefly, 30 min after mixing with 2-hour-incubated spermatozoa, cumulus-free eggs were fixed with 0.25% glutaraldehyde. The bound spermatozoa were observed with an Olympus IX73 microscope.

### Immunostaining of mouse sperm

Immunostaining of mouse sperm was performed as described previously (Yamaguchi et al., 2008). Briefly, all samples were mounted on glass slides and dried. After washing with PBS, slides were blocked with 10% Normal Goat Serum/ PBS (B6J) or 10% Newborn Calf Serum (NBCS)/ PBS (BDF1) for 1 hour and incubated with primary antibodies (rat monoclonal anti-mouse SPACA4 (KS139-281) (Fujihara et al., 2013b); rat monoclonal anti-mouse IZUMO1 (KS64-125) (Fujihara et al., 2013b; Ikawa et al., 2011); rabbit anti-mouse IZUMO1 (Inoue et al., 2005)) in blocking buffer at 4°C overnight. After washing with PBS-T (B6J) or 10% NBCS/PBS-T (BDF1), the slides were incubated with secondary antibodies (goat anti-rat IgG Alexa Fluor 488 (A11006); goat anti-rat IgG Alexa Fluor 546 (A11081); goat anti-rabbit IgG Alexa Fluor 488 (A11008) (all purchased from Thermo Fisher Scientific)) in blocking buffer for 1 hour. After washing with PBS containing 0.05% Tween-20, the slides were observed under a fluorescence microscope (B6J: Axio Imager.Z2, Zeiss; BDF1: IX70, Olympus). The mouse strain *Tg(Acr-EGFP)1Osb* was used to identify spermatozoa with unreacted acrosomes (Nakanishi et al., 1999).

### Western blotting

Immunoblot analysis from mouse sperm was performed as described previously (Yamaguchi et al., 2006). Briefly, sperm samples were collected from the cauda epididymis and vas deferens. These samples were homogenized in lysis buffer containing 1% Triton X-100 and 1% protease inhibitors (Nacalai Tesque), centrifuged, and the supernatants were collected. Protein lysates were resolved by SDS/PAGE under reducing condition and transferred to PVDF membranes (Merck Millipore). After blocking, blots were incubated with primary antibodies (rat monoclonal antimouse SPACA4 (KS139-281) (Fujihara et al., 2013b); goat polyclonal anti-BASIGIN sc-9757 (Santa Cruz Biotechnology)) overnight at 4°C, and then incubated with secondary antibodies conjugated with horseradish-peroxidase (HRP) (HRP-conjugatedgo at anti-rat immunoglobulins (IgGs) (112-035-167) and HRP-conjugated goat anti-mouse IgGs (115-036-062) (Jackson ImmunoResearch Laboratories)). The detection was performed using an ECL plus Western blotting detection kit (GE Healthcare).

### Protein sequence alignments of fertility factors

Protein sequences of human CD9 (P21926), IZUMO1 (Q8IYV9), SPACA6 (W5XKT8), JUNO/ IZUMO1R (A6ND01), TMEM95 (Q3KNT9), FIMP (Q96LL3) and mouse LLCFC/SOF1 (Q9D9P8) were downloaded from uniprot (https://www.uniprot.org/), and searched via NCBI protein BLAST blastp (https://blast.ncbi.nlm.nih.gov/Blast.cgi?PAGE=Proteins) for homologous protein sequences in *Mus musculus* (taxid:10090) or *Homo sapiens* (taxid:9606), reptiles (taxid:8459), Xenopus (taxid:8353), fugu (taxid:31032), *Danio rerio* (taxid:7955), platypus (taxid:9258) and armadillo (taxid:9359). If no homologous protein sequence was found, the search was extended to translated nucleotide sequences using tblastn. Protein sequence alignments and percent identity matrices were generated with clustal omega (https://www.ebi.ac.uk/Tools/msa/clustalo/) using default parameters and plotted with JalView (https://www.jalview.org/).

### Bioinformatic analyses of Ly6/uPAR genes

Our analysis of the mouse Ly6/uPAR family considered 66 mouse genes annotated in the Ensembl release 79 of GRCm38. The Ensembl release was chosen to match the mouse expression data (Li et al., 2017). The set of genes was selected based on sequence similarity to the mouse Ly6/ uPAR family published in (Loughner et al., 2016) and domain similarity searches with Ly6/uPAR domain HMMs against the Ensembl proteome (Herberg et al., 2018). For defining the set of human orthologs of the mouse Ly6/uPAR genes, mouse gene symbols were used to query against Diopt v8.0 (Hu et al., 2011), and hits were only considered with Best.Score, Best.Score.Reverse and Rank “high”. Mouse expression data were obtained from (Li et al., 2017). Human expression data were obtained from GTEx v8 (www.gtexportal.org; GTEx_Analysis_2017-06-05_v8_RNASeQCv1.1.9_gene_media_n_tpm). *Xenopus laevis* expression data were derived from the NCBI GEO entry GSE73419 (Session et al., 2016). *Danio rerio* expression data were derived from GEO entries GSE111882 (testis, ovary, mature oocytes) (Herberg et al., 2018), GSE147112 (oogenesis, mature oocytes) (Cabrera-Quio et al., 2021) and GSE171906 (adult tissues) (Noda et al., 2021). Heatmaps were used to illustrate the relative expression levels of genes across tissues. To this end, published library size and length corrected expression values (TPM/ FPKM) were obtained; after square-root-based variance stabilizing transformation the obtained values were centered and scaled per gene (z-score).

### Statistical analysis

Statistical analysis was performed with the GraphPad Prism 7 software. Statistical tests are detailed in each Figure legend. Differences were considered significant at p < 0.05 (*) (p < 0.01 (**); p < 0.001 (***); n.s., not significant). Error bars represent standard deviation. Figure legends indicate the number of *n* values for each analysis.

## Data availability statement

All data are available in the manuscript or the supplementary material and are available from the corresponding authors upon request.

## List of primers

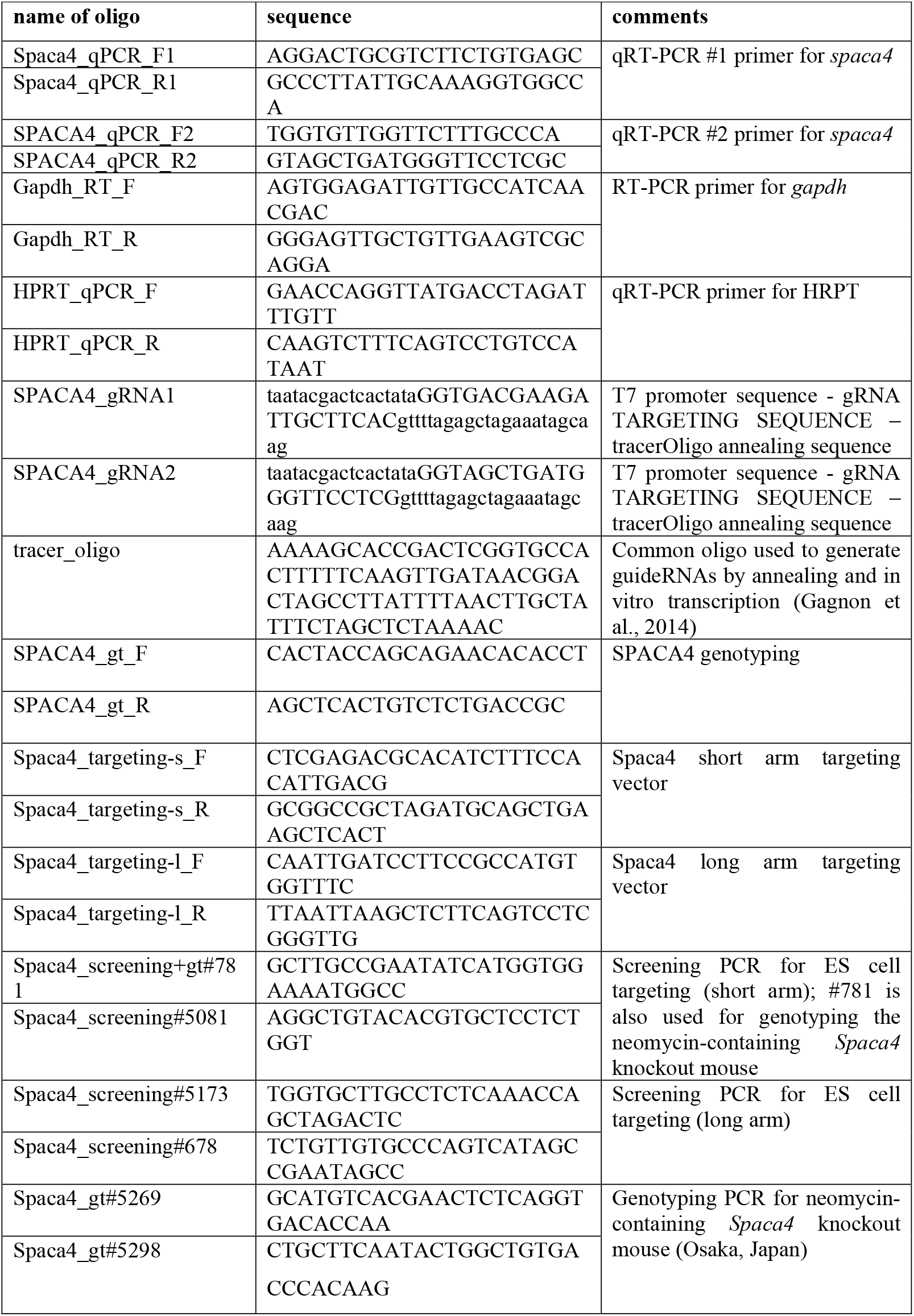

## SUPPLEMENTARY FIGURES

### Supplementary Figure 1

**Fig S1.**
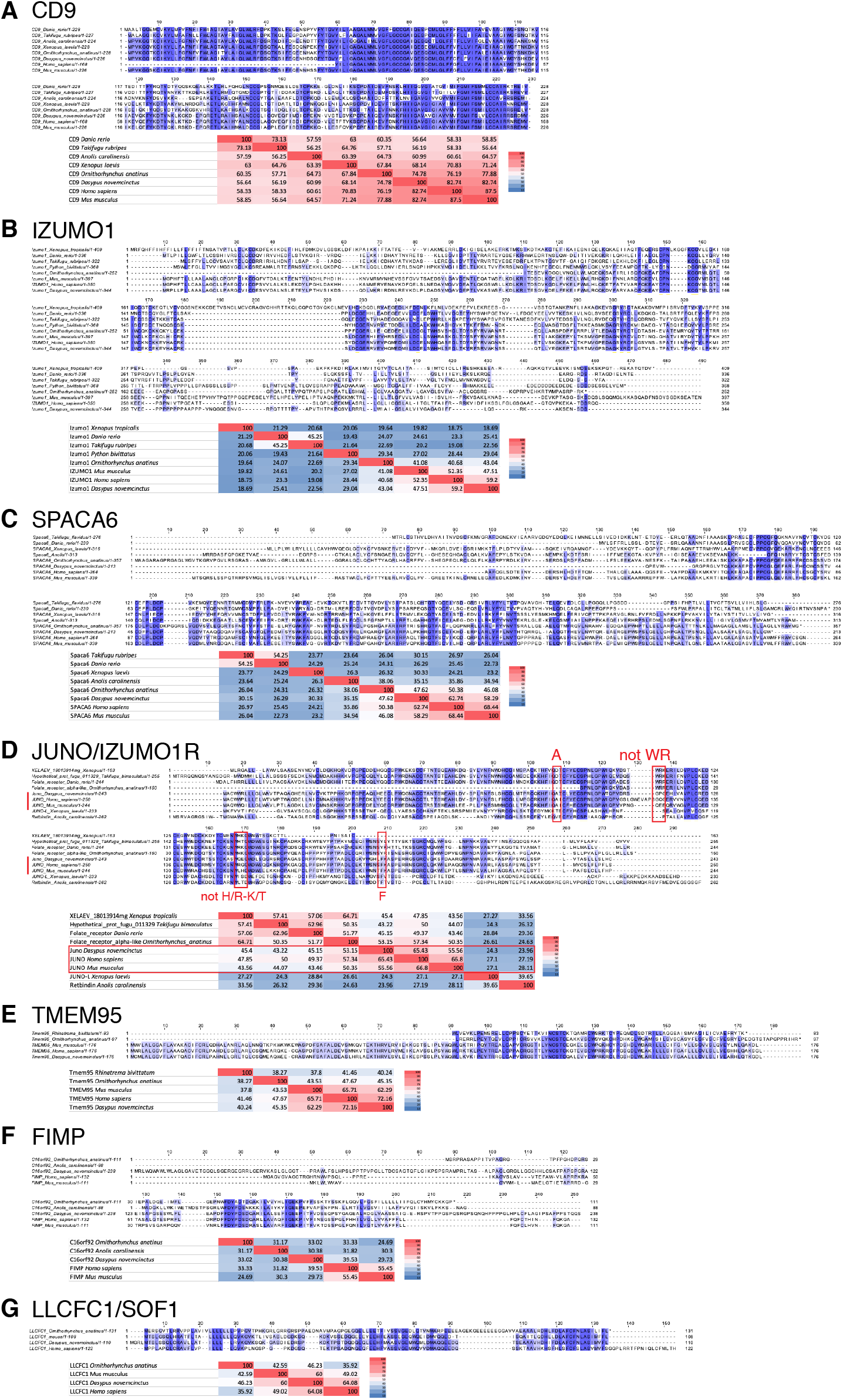
Analysis of the conservation of mammalian fertility factors across vertebrates. (**A-G**) **Protein sequence alignments** (top) and **percent identity matrices** (bottom) **of mammalian fertility factors known to be essential for sperm-egg interaction in mice**. (**A**) CD9, (**B**) IZUMO1, (**C**) SPACA6, (**D**) JUNO/IZUMO1R, (**E**) TMEM95, (**F**) FIMP, (**G**) LLCFC1/SOF1. To **D**: Although folate receptor homologs (JUNO/IZUMO1R belong to the folate receptor family) are found outside of mammals, amino acids defining the JUNO/IZUMO1R (highlighted in red) are different outside of mammals (Grayson, 2015). Based on this definition, JUNO/IZUMO1R homologs are only present in mammals (species highlighted in the red box in the percent identity matrix).

### Supplementary Figure 2

**Fig S2.**
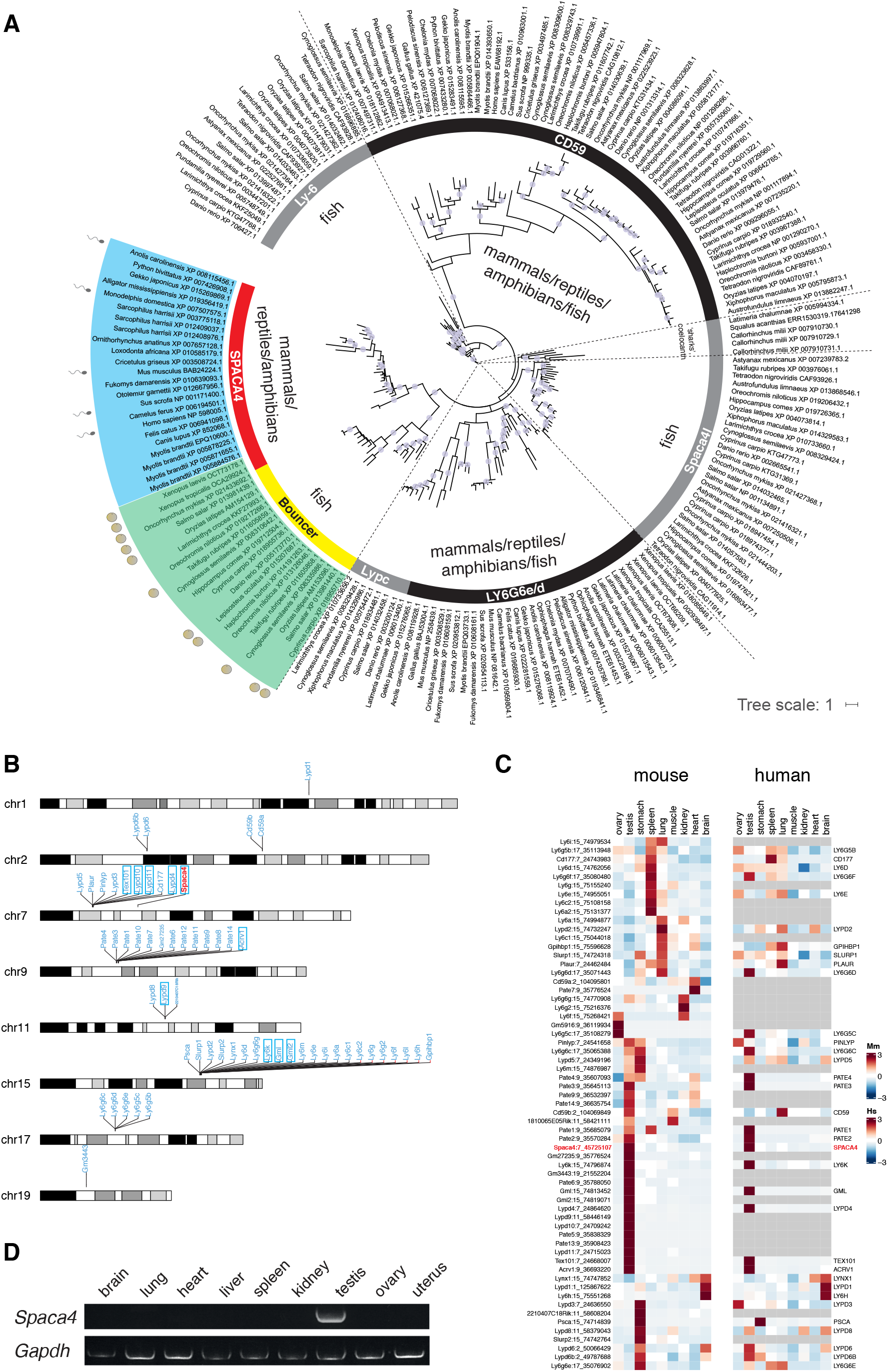
Expression analysis of mammalian Ly6/uPAR proteins. (**A**) **Phylogenetic tree of selected groups of Ly6/uPAR proteins across vertebrates**. Maximum-likelihood phylogenetic tree based on Ly6/uPAR protein sequence alignments across vertebrates (adapted from (Herberg et al., 2018)). Branches supported by ultrafast bootstrap values (>=95%) are marked with a blue dot. SPACA4 (red) is present in mammals, reptiles and amphibians; Bouncer (yellow) is present in fish. The sperm and egg symbols next to selected species’ genes indicate expression in testis (blue) or ovaries (green). (**B**) **Genomic localization of Ly6/uPAR genes in mice.** Ly6/uPAR genes are clustered in the genome. Only chromosomes containing at least one Ly6/uPAR gene are shown. Testis-expressed genes are highlighted with a blue box; *Spaca4* is highlighted in red. (**C**) **Mouse and human Ly6/uPAR proteins are expressed in various tissues, including in gametes** (extension of **Fig 1B** including human orthologs). Heatmaps of expression levels of homologous mouse (Mm; left) and human (Hs; right) Ly6/uPAR proteins (homologous proteins are shown in one row) across various adult tissues. Heatmaps are color-coded based on z-scores of the normalized gene expression values (average of the squareroot) of RNA-Seq data from murine and human adult tissues (Li et al., 2017) (www.gtexportal.org). Adult tissues are indicated at the bottom. Gene names are given on the left (mouse) and right (human) of the corresponding heat-maps. Numbers behind gene names indicate chromosome locations in mice or humans. *Spaca4* is highlighted in red. (**D**) **Murine *Spaca4* is expressed in the male but not the female germline.** RT-PCR for *Spaca4* from cDNA of the indicated mouse tissues relative to the housekeeping gene *Gapdh*. The template cDNA was generated from a mixture of the tissue from 3-4 different mice.

**Fig S3.**
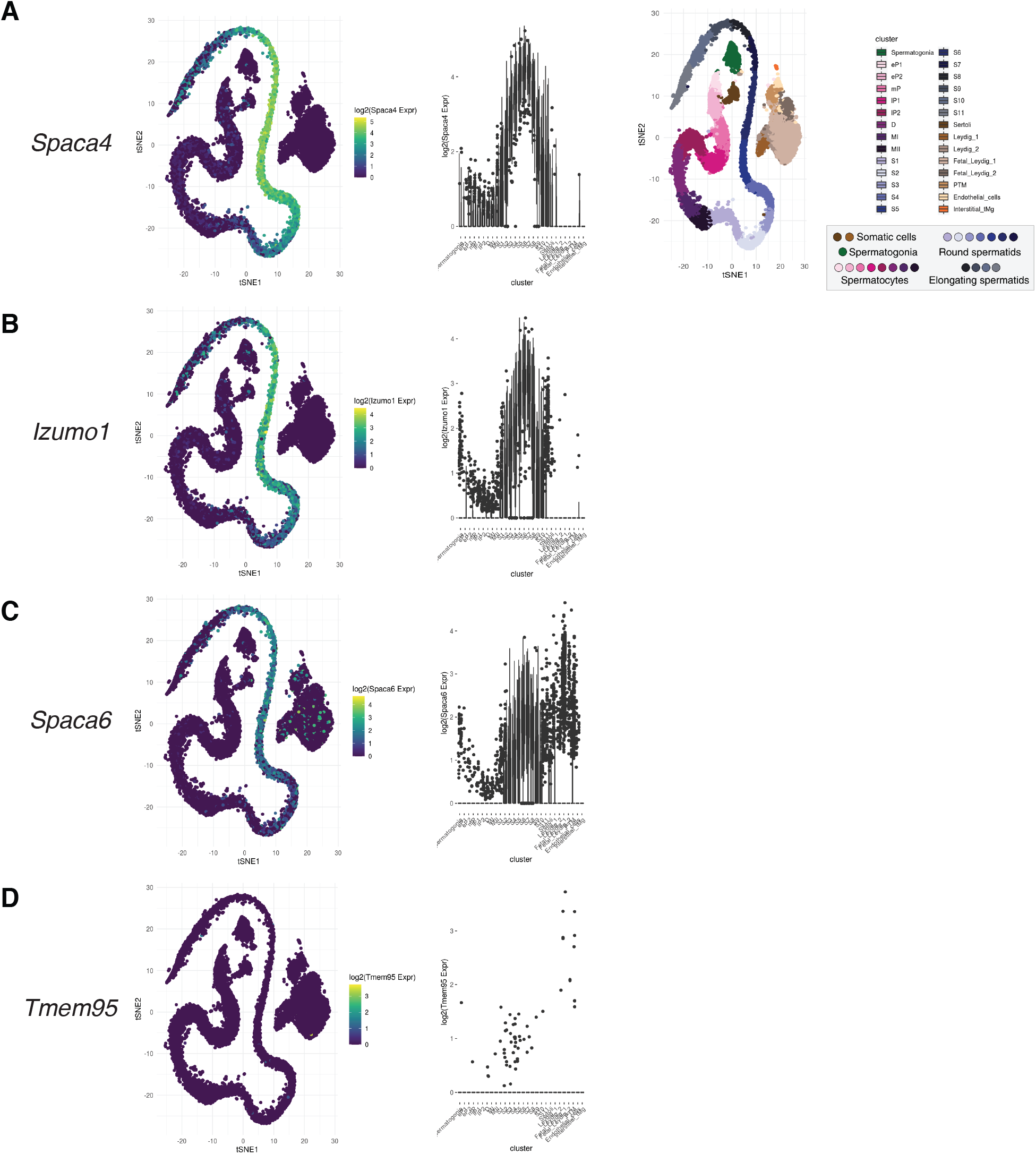
Analysis of the expression of selected mammalian fertility factors during murine spermatogenesis. (**A-D**) Expression values for selected fertility factors during murine spermatogenesis were derived from published single-cell RNA-Seq data (Ernst et al., 2019). The web-browser version https://marionilab.cruk.cam.ac.uk/SpermatoShiny/ was used to generate the plots for (A) *Spaca4*, (B) *Izumo1*, (C) *Spaca6*, (D) *Tmem95*. The *Spaca4* temporal expression pattern resembles *Izumo1’s* pattern.

**Fig S4.**
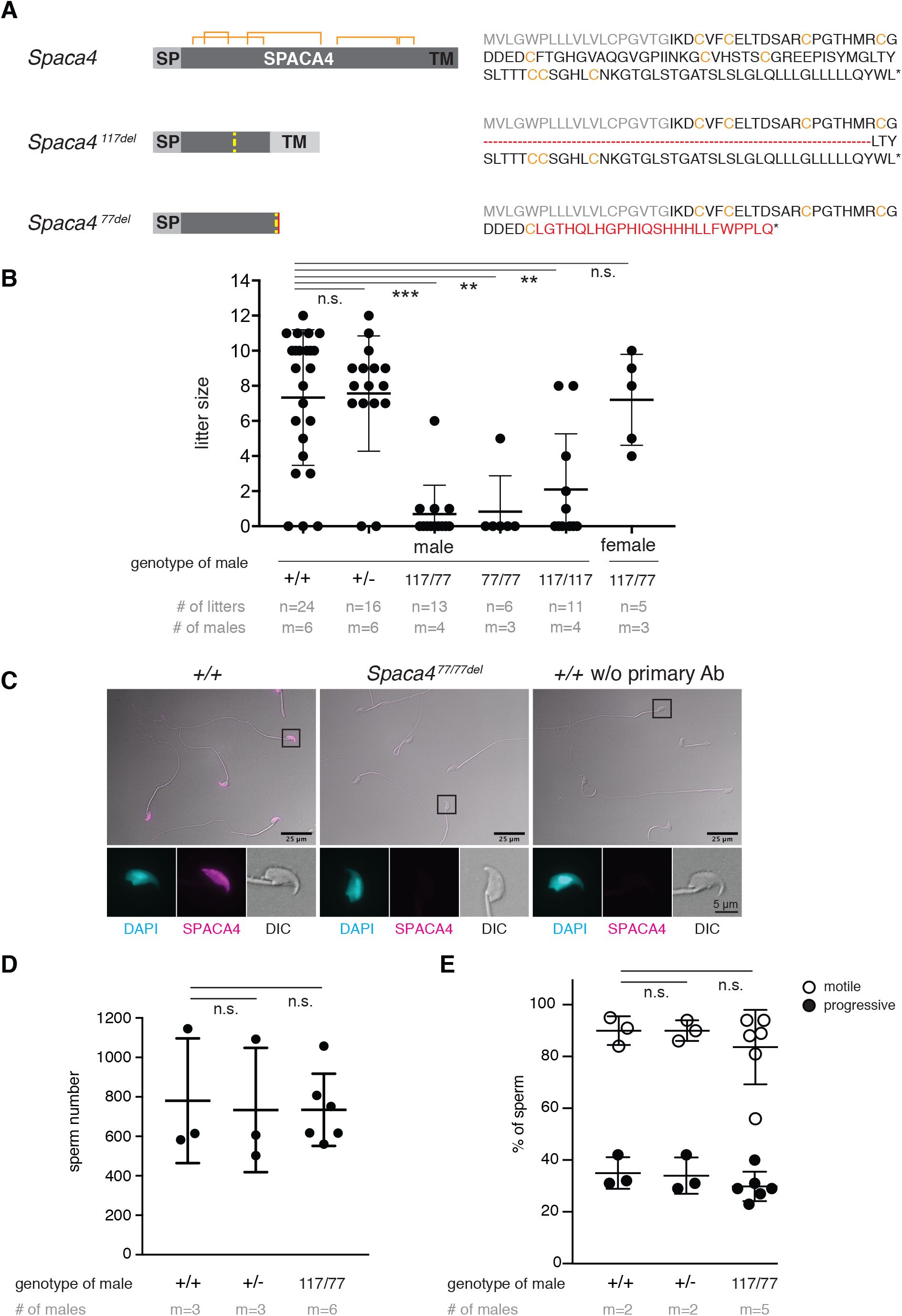
Generation and phenotypic analysis of Spaca4 knockout mice. (**A**) **Overview of the *C57BL/6J-Spaca4* knockout alleles generated by CRISPR/Cas9-mediated targeted mutagenesis**. (Left) Schematics of the wild-type and knockout alleles. Yellow dashed lines indicate the site of the deletions. Predicted disulfide bridges are indicated in orange. SP, signal peptide; TM, transmembrane region. (Right) Amino acid sequences of the protein products that are predicted to be produced from the different alleles. One mutant allele contains a 117-nt in-frame deletion after amino acid 42. The other allele contains a 77-nt out-of-frame deletion after amino acid 47. Cysteines predicted to form disulfide bridges are indicated in orange; the SP sequence is shown in grey; the out-of-frame additional protein sequence in the 77del allele is shown in red. (**B**) ***Spaca4* knockout male mice are sub-fertile**. Litter sizes of C57BL/6J-*Spaca4* wild-type (+/+), heterozygous (+/−), transheterozygous (117/77) and homozygous (77/77 or 117/117) males of the indicated allele combinations caged with B6129F1 wild-type females, or B6129F1 wild-type males caged with transheterozygous (−/−) females. Successful mating was confirmed by plug checks. Data are means ± SD. ***p < 0.0001, **p < 0.001 (Kruskal-Wallis test with Dunn multiple-comparisons test); n.s., not significant. n = number of litters; m = number of male mice tested. (**C**) **SPACA4 protein is absent in morphologically normal sperm of *Spaca4* knockout mice**. Immunostaining of sperm detects SPACA4 protein (magenta) under permeabilizing conditions in the head of sperm derived from wild-type (*+/+*) but not *Spaca4* knockout (77/77del) mice. DAPI (cyan) staining labels the sperm nucleus. A control immunostaining in which the primary antibody was omitted is shown on the right. DIC, differential interference contrast image. Scale bar, 25 μm. (**D**) **Sperm number is normal in *Spaca4* knockout males**. Sperm number of *Spaca4* wild-type (+/+), heterozygous (+/−) and transheterozygous (−/−) males. Data are means ± SD. n.s., not significant (one-way ANOVA with Dunnett’s multiple-comparisons test). m = number of male mice tested. (**E**) **Overall sperm motility is not affected in *Spaca4* knockout males.** Sperm motility of *Spaca4* wild-type (+/ +), heterozygous (+/−) and transheterozygous (−/−) males. Shown is the percentage of sperm that was motile (open circles, comparison +/+ versus −/−: p = 0.65) and progressively motile (closed circles, comparison +/+ versus −/−: p = 0.41). Data are means ± SD. n.s., not significant (One-way ANOVA with Dunnett’s multiple-comparisons test). m = number of male mice tested.

**Fig S5.**
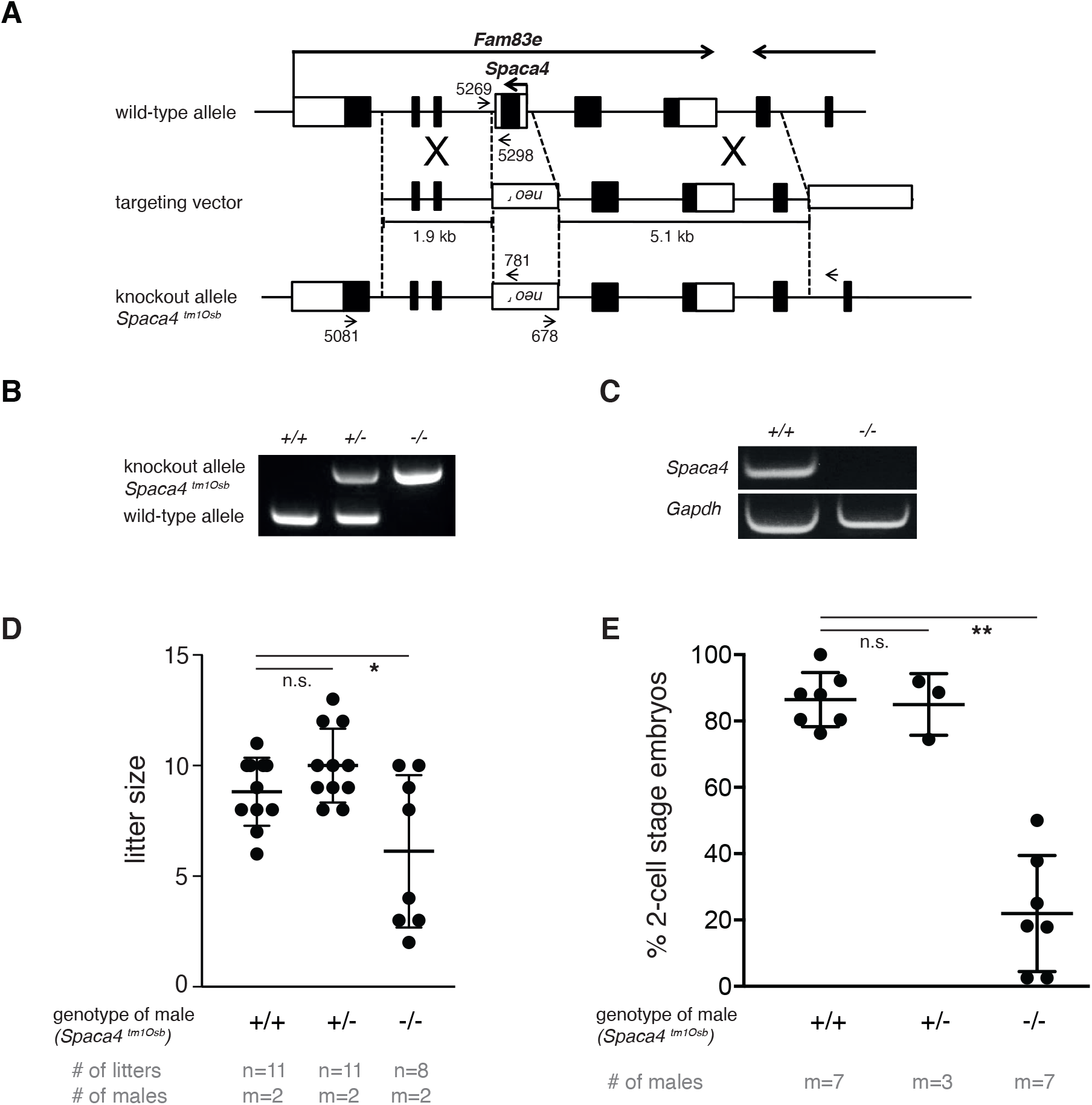
Male *Spaca4* knockout mice are sub-fertile. (**A**) **Targeted disruption of the murine *Spaca4* gene to generate *Spaca4*^tm1Osb^**. To disrupt the *Spaca4* gene, the single exon was replaced with a neomycin resistance cassette (*neo*), and a thymidine kinase cassette (*tk*) was used for negative selection. Labeled arrows indicate primer binding sites. (**B**) **Genotyping of *Spaca4*^tm1Osb^ knockout mice**. Both the wild-type allele (a 0.3-kb band) and the knockout allele (a 0.5-kb band) were amplified by PCR, using primers #5269 and #5298 for the wild-type allele and primers #781 and #5269 for the knockout allele. (**C**) **RT-PCR analysis of testis in wild-type and *Spaca4*^tm1Osb^ knockout mice**. The *Spaca4*-specific 236-nt band was amplified from wild-type but not from *Spaca4*^tm1Osb^ knockout testis. *Gapdh* was used as a control. **(D-E**) ***Spaca4*^tm1Osb^ knockout mice are sub-fertile**. (**D**) *Spaca4*^tm1Osb^ knockout males (−/−) copulated normally but were sub-fertile compared to wild-type (+/+) and heterozygous *Spaca4*^tm1Osb^ (+/−) mutant males (comparison +/+ versus −/−: p = 0.0333, unpaired t-test). Data are means ± SD. n.s., not significant. n = number of litters; m = number of male mice tested. (**E**) *In vitro* fertilization of oocytes from wild-type females using sperm from wildtype (+/+), heterozygous *Spaca4*^tm1Osb^ (+/−) or homozygous *Spaca4*^tm1Osb^ (−/−) males. Plotted is the percentage of 2-cell stage embryos as a measure of successful fertilization. Data are means ± SD. p = 0.003 (Kruskal-Wallis test with Dunn multiple-comparisons test); n.s., not significant. m = number of males tested.

**Fig S6.**
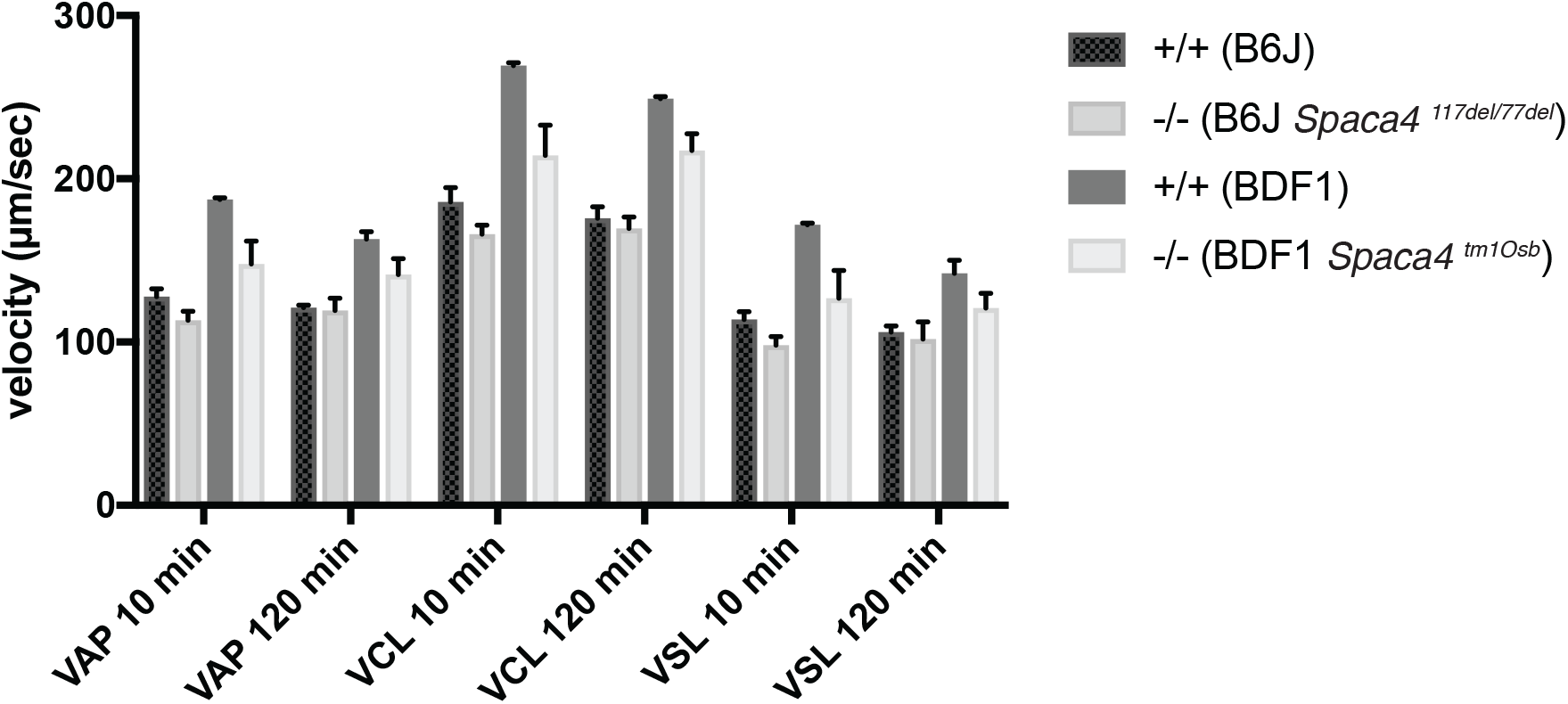
*Spaca4* knockout sperm has reduced motility under IVF conditions. Sperm motility was assessed under IVF conditions using the computer-assisted sperm analysis system CEROS II. Sperm motility of transheterozygous B6J (117del/77del; −/−) males was compared to wild-type B6J (+/+) males; similarly, sperm motility of BDF1 *Spaca4*^tm1Osb^ knockout (−/−) males was compared to wildtype BDF1 (+/+) males to control for possible differences in the genetic backgrounds of both mutants. There are significant differences of all three parameters (VAP, VCL, and VSL) between wild-type and *Spaca4* mutant (transheterozygous B6J and BDF1 knockout) spermatozoa after 10 min incubation (p < 0.05). After 120 min incubation, however, there are no significant differences between B6J backgrounds although there are significant differences between BDF1 backgrounds. Plotted is the velocity in μm/sec. VAP: average path velocity, VSL: straight line velocity, VCL: curvilinear velocity. Data are means ± SD. n.s., not significant (Student’s t test). m = 3 male mice tested for each genotype.

